# A New Model of SARS-CoV-2 Infection Based on (Hydroxy) Chloroquine Activity

**DOI:** 10.1101/2020.08.02.232892

**Authors:** Robert J. Sheaff

**Author notes:** **Materials & Correspondence** Robert J. Sheaff, Associate Professor, The University of Tulsa, Department of Chemistry and Biochemistry, Keplinger M2215, 800 South Tucker Drive, Tulsa, OK 74104, Office.

## Abstract

Chloroquine and hydroxychloroquine (H)CQ are well known anti-malarial drugs, while their use against COVID-19 is more controversial. (H)CQ activity was examined in tissue culture cells to determine if their anti-viral benefits or adverse effects might be due to altering host cell pathways. Metabolic analysis revealed (H)CQ inhibit oxidative phosphorylation in mitochondria, likely by sequestering protons needed to drive ATP synthase. This activity could cause cardiotoxicity because heart muscle relies on beta oxidation of fatty acids. However, it might also explain their therapeutic benefit against COVID-19. A new model of SARS-CoV-2 infection postulates virus enters host cell mitochondria and uses its protons for genome release. Oxidative phosphorylation is eventually compromised, so glycolysis is upregulated to maintain ATP levels. (H)CQ could prevent viral infection and/or slow its replication by sequestering these protons. In support of this model other potential COVID-19 therapeutics also targeted mitochondria, as did tobacco smoke, which may underlie smokers’ protection. The mitochondria of young people are naturally more adaptable and resilient, providing a rationale for their resistance to disease progression. Conversely, obesity and diabetes could exacerbate disease severity by providing extra glucose to infected cells dependent on glycolysis. The description of (H)CQ function presented here, together with its implications for understanding SARS-CO-V2 infection, makes testable predictions about disease progression and identifies new approaches for treating COVID-19.

## Introduction

Severe Acute Respiratory Syndrome Coronavirus-2 (SARS-CoV-2) is the causative agent for COVID-19, a disease that has killed hundreds of thousands and severely impacted the world economy.^1,2^ While much is known about its structure and replication, effective treatments remain elusive. The coronavirus is so named due to spike-like glycoproteins on its surface that bind the angiotensin-converting enzyme (ACE-2) receptor, allowing virus to fuse with the cell membrane and enter as an endosome.^3,4^ The positive strand RNA genome is thought to be released into the host cell after endosome acidification and dissociation of associated capsid proteins.^5^ This RNA serves as both mRNA for production of viral proteins and template for synthesizing additional genomic copies.^5^ New viral particles are assembled and released via exocytosis.^6^ Anti-viral drugs can target unique viral processes or disrupted host systems to prevent virus proliferation, cause host cell death, or illicit an immune response. ^7^, (H)CQ are well characterized anti-malarial drugs used extensively over the past 60 years.^8^ Some clinical studies suggest they act as preventive and therapeutic agents against COVID-19, while others adamantly dispute these claims.^9–11^ Proposed mechanism(s) of action are also controversial. One hypothesis is based on their anti-malarial activity, positing that (H)CQ protonation and accumulation in the endosome prevents the acidification required for genome release.^12^ Alternatively, it has been proposed that (H)CQ act as ionophoric agents that transport zinc ions into infected cells, where they inhibit the viral RNA replicase enzyme.^13^ (H)CQ can also dampen excess immune responses (e.g. in lupus) and so they might mitigate the hyperactive immune response sometimes associated with COVID-19.^14,15^ Both CQ and HCQ have detrimental side effects (e.g. cardiotoxicity) that raise further questions about their therapeutic value.^16^ In view of these controversies (H)CQ effects on human tissue culture cells was investigated, in hopes that clarifying their mechanism of action will lead to a better understanding of SARS-Co-V2 infection and help identify more effective therapies with reduced side effects.

## Results

### (H)CQ inhibits mitochondrial ATP production

Treating h293 cells with CQ overnight had little effect on cell viability (resazurin based assay), but caused a concentration dependent decrease in ATP levels (luciferase based assay) (Figure 1A). A control experiment showed CQ was not simply inhibiting luciferase activity (Supl Fig. 1). While CQ appeared to target cell metabolism, it was unclear whether cytosolic glycolysis or mitochondrial oxidative phosphorylation was affected. To address this question cells were treated for 2hrs with well-defined metabolic inhibitors to determine which pathway(s) actively produce ATP (Figure 1B). Surprisingly, neither the glucose analog 2-deoxy-glucose (2DG) nor the Electron Transport Chain (ETC) inhibitor rotenone decreased ATP levels. When combined, however, they acted synergistically to completely block ATP production. These results indicate that when either glycolysis or oxidative phosphorylation is blocked cells maintain ATP levels using the uninhibited pathway. Since 2DG inhibits glycolysis (and hence pyruvate formation), cells are likely relying on amino acid metabolism via the TCA/ETC. Consistent with this interpretation rotenone alone decreased ATP levels when amino acids were the only carbon source (Figure 1C). As expected, adding back glucose re-established glycolysis and overcame rotenone inhibition.

**Figure 1:**
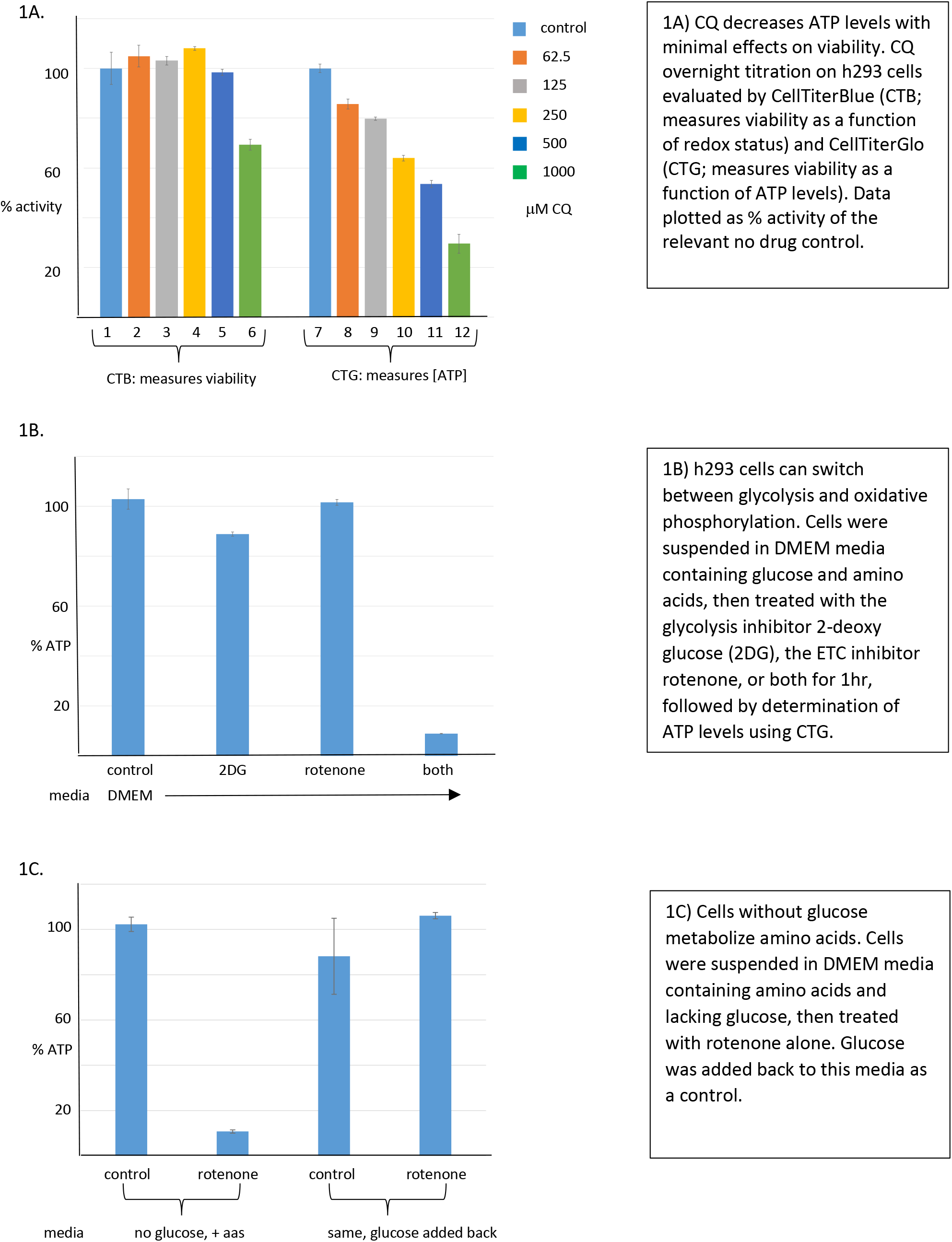
Chloroquine inhibits ATP production.

(H)CQ effects on specific metabolic pathways could now be examined by pre-treating cells with 2DG or rotenone (Figure 2A). A short 2hr CQ exposure had no effect on ATP levels when both glycolytic and oxidative phosphorylation pathways were available (lanes 1-6). When cells were pre-treated with rotenone to inhibit the ETC and force a switch to glycolysis, CQ again had no effect (lanes 13-18). However, when glycolysis was inhibited with 2DG and cells utilized oxidative phosphorylation, CQ decreased ATP levels in a concentration dependent manner (lanes 7-12). Similar data were obtained with HCQ (Figure 2B). These results were repeated in immortalized but untransformed human diploid fibroblasts (Supl. Fig. 2), indicating that this (H)CQ activity is not confined to h293 cells. Taken together these data indicate (H)CQ is specifically inhibiting mitochondrial ATP production. In support of this idea CQ alone reduced ATP levels when cells were forced to metabolize amino acids (Figure 2C, lanes 1-6), while adding back glucose overcame this inhibition by allowing glycolysis (lanes 7-12). An alternative explanation is that CQ inhibits an early step in amino acid metabolism. To rule out this possibility the experiment was repeated using L-15 media, which substitutes galactose for glucose. Galactose is metabolized via the glycolytic pathway, but because this process costs an additional 2 ATP, the resulting pyruvate must be metabolized in mitochondria to generate net ATP.^17^ Note: this media must be supplemented with sodium bicarbonate, otherwise its rapid acidification in a CO_2_ incubator blocks CQ activity (Supl Fig.3, and described in more detail below). Control experiments using 2DG and rotenone confirmed cells in L-15 media were metabolizing galactose (Supl. Fig. 4). CQ and HCQ alone inhibited ATP production under these conditions (Figure 2D, lanes 1-6 and 13-18), while adding back glucose activated glycolysis and overcame inhibition (lanes 7-12 and 19-24). Thus, (H)CQ specifically targets mitochondrial oxidative phosphorylation rather than amino acid metabolism. Given the well described ability of (H)CQ to be protonated and trapped in acidic compartments, CQ likely inhibits ATP production by depleting protons in the mitochondrial intermembrane space needed to run ATP synthase.^15^

**Figure 2:**
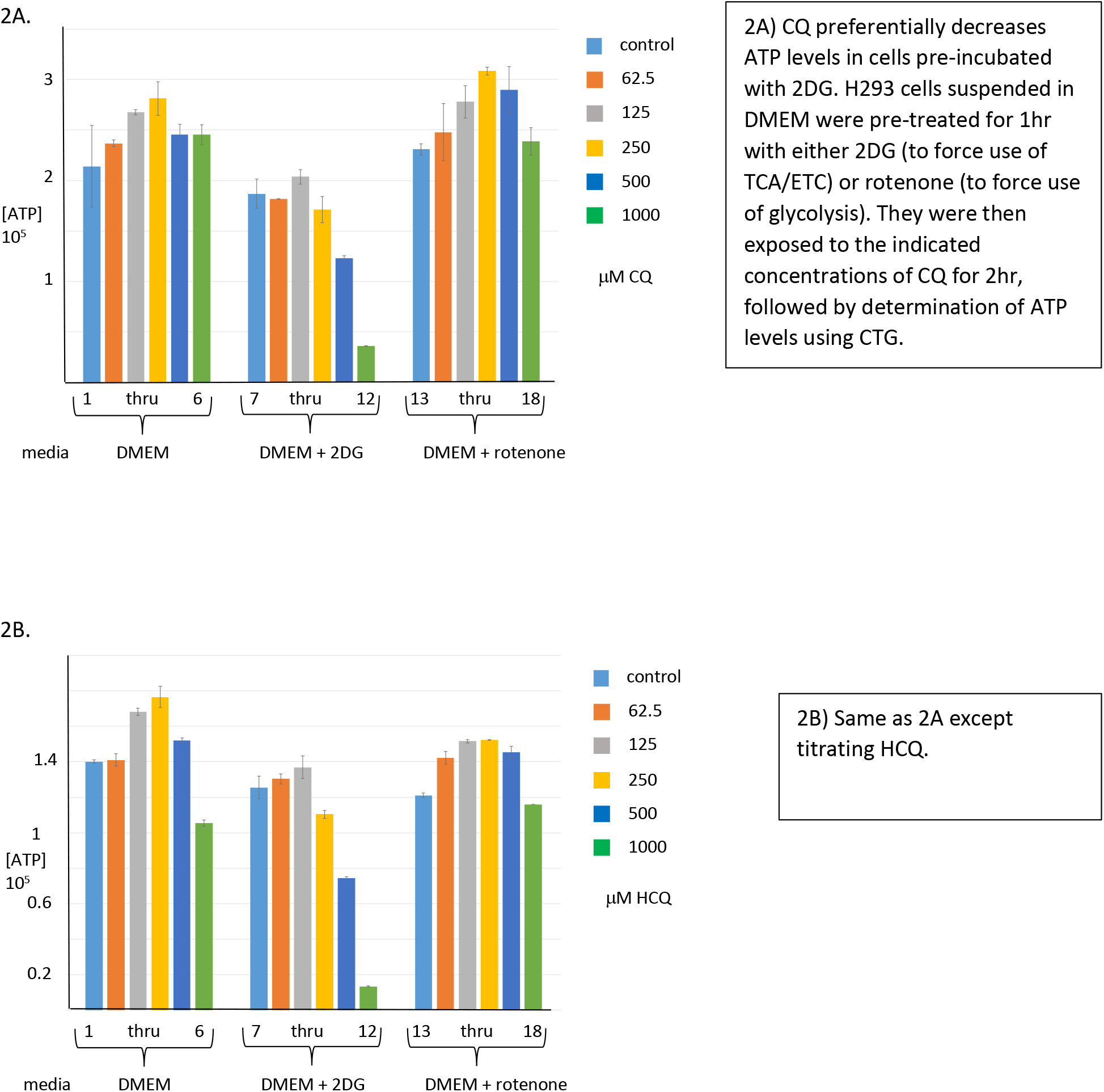
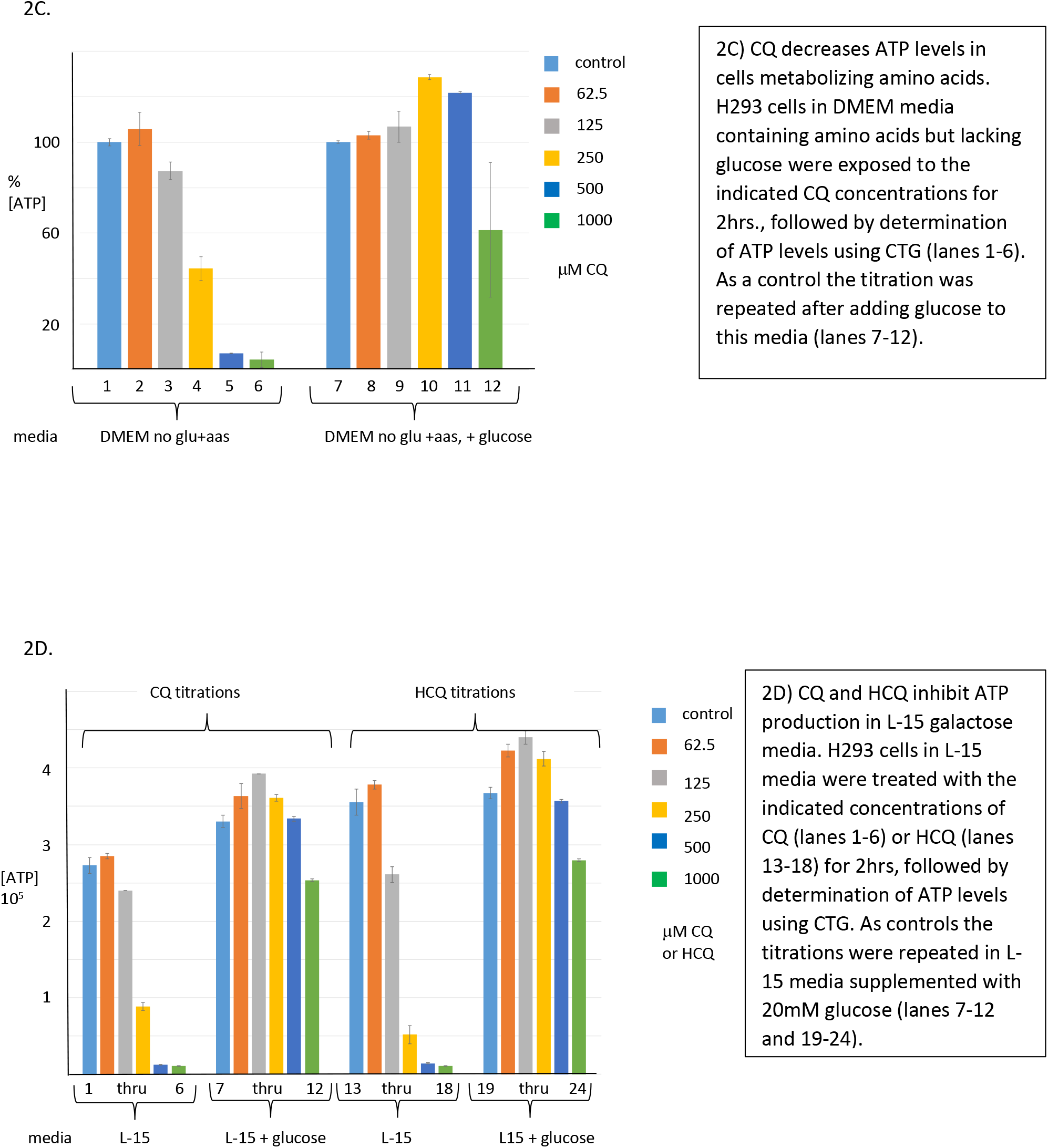
Chloroquine inhibits mitochondrial ATP production.

### (H)CQ activity is time and pH dependent

The above hypothesis implies that only the free base (unprotonated) form of (H)CQ enters cells, where it is then protonated and trapped. Given that the media pH is ~ 7.4 and the pKas for CQ nitrogens are above that (8.3 and 10.1), CQ exists mainly in the protonated form when added to cells (Figure 3A). Thus, cell entry is likely to be the rate limiting step. To test this idea cells without and with CQ were pre-incubated for various times, washed and pelleted, then re-suspended in L-15 media followed by determination of ATP levels (Figure 3B). As predicted the amount of inhibition increased with pre-incubation time. Rate limiting cell entry was tested more directly by varying media pH. As media basicity increased (generating more of the free base form) CQ inhibition also increased, consistent with its equilibrium determining rate of cell entry (Figure 3C). This result explains why CQ failed to inhibit in L-15 media lacking sodium bicarbonate (Supl. Fig. 3).

**Figure 3:**
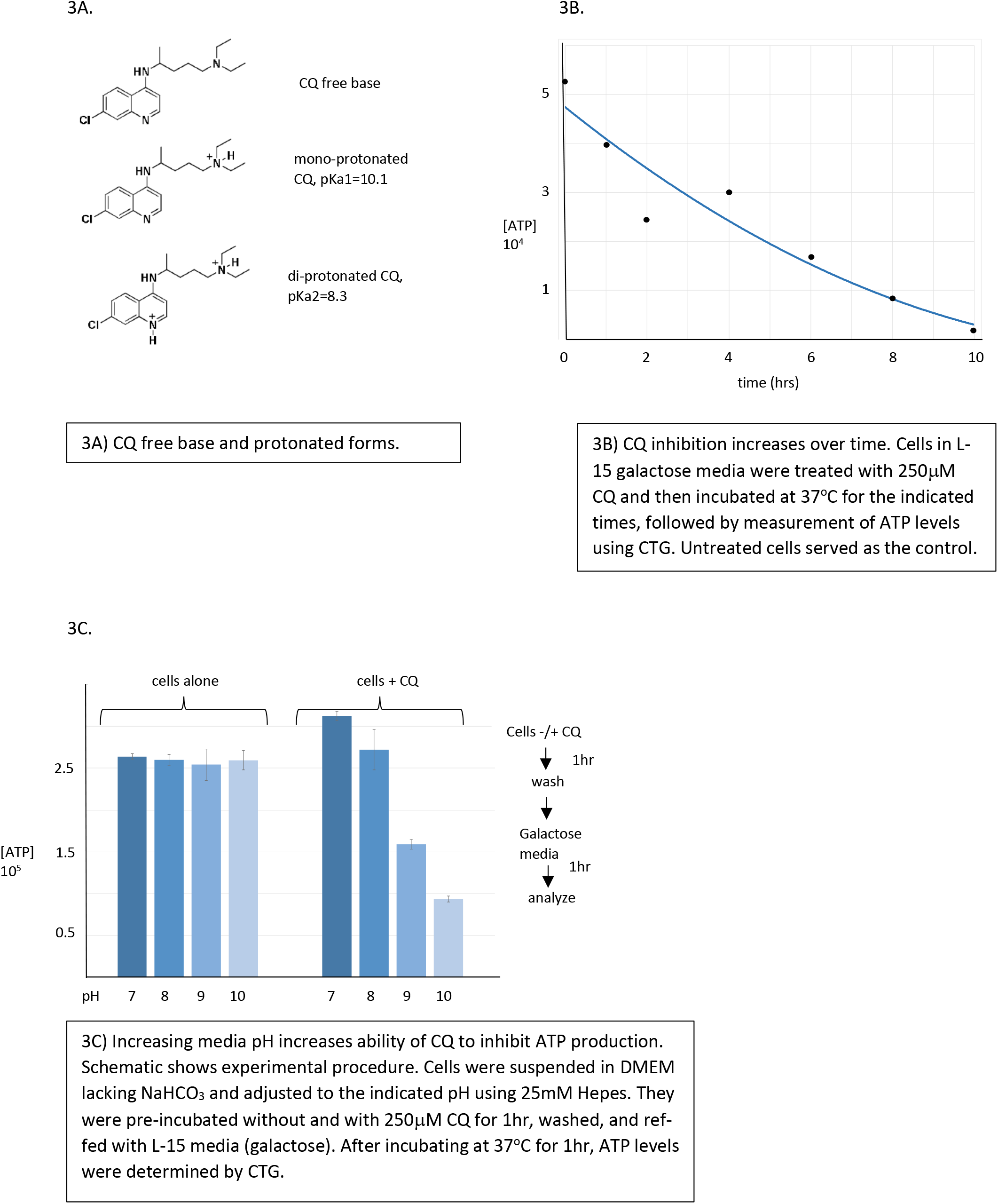
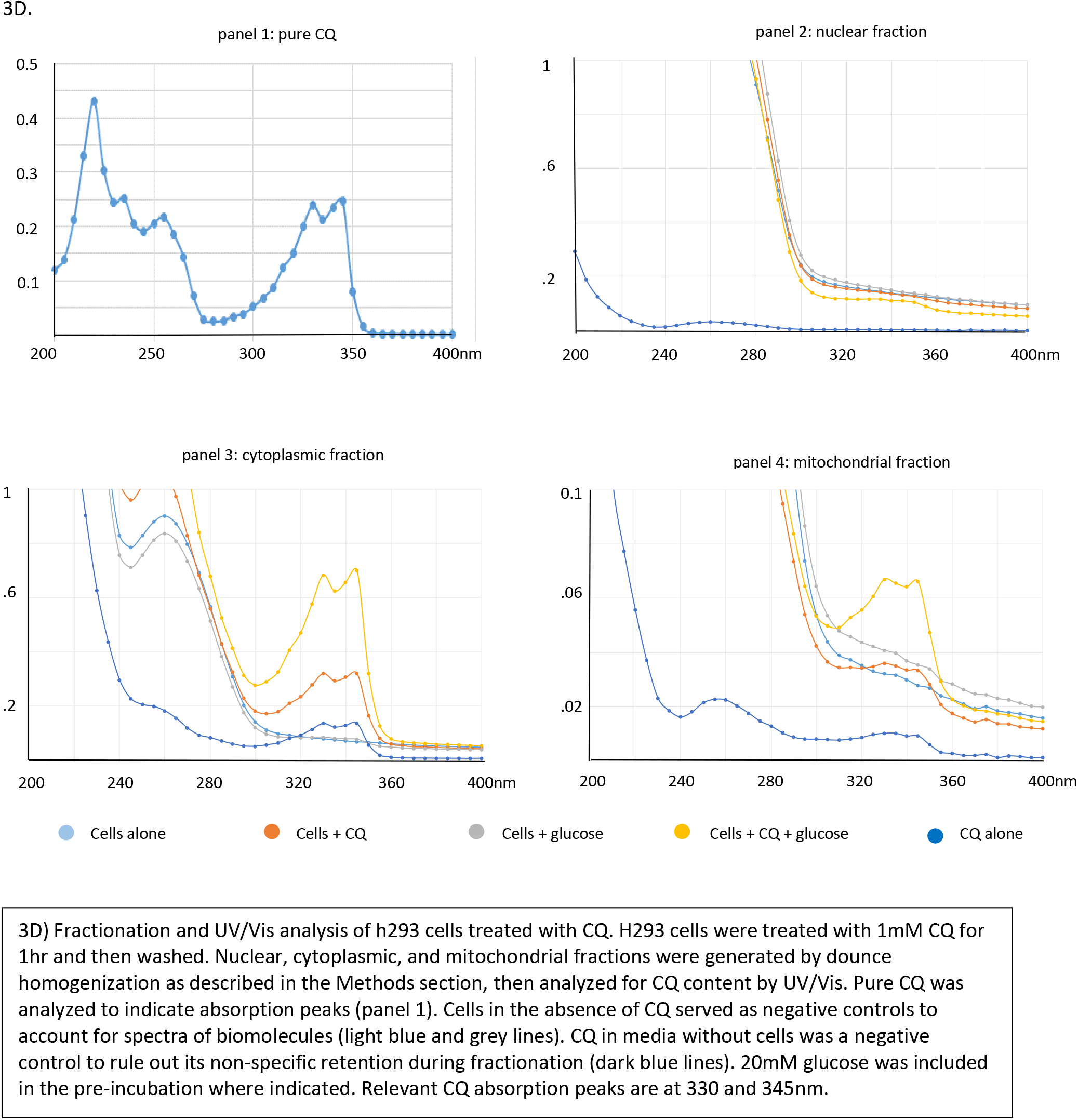
CQ entry into cells and inhibition of mitochondrial ATP production is time and pH dependent.

Cell fractionation experiments and UV/Vis spectroscopy were performed to further confirm cell entry and identify CQ location(s) (Figure 3D). Pure CQ has two distinctive peaks at ~ 330 and 345nm (panel 1) that can be used for identification. After fractionating cells incubated with CQ by dounce homogenization, the drug was observed in the cytoplasmic (panel 3) but not nuclear fraction (panel 2). Including glucose during the pre-incubation increased the amount of cytoplasmic CQ, which may reflect its protonation in this compartment due to lactic acid produced by glycolysis. This supposition is supported by a similar result in red blood cells, which lack both a nucleus and mitochondria (Supl. Fig. 5). Finally, the mitochondrial fraction was examined. There was modest CQ accumulation in mitochondria (panel 4, orange line) that increased dramatically when glucose was present during the pre-incubation (panel 4, yellow line). Since glucose also elevated cytoplasmic CQ (panel 3), it likely increased the probability of it eventually becoming trapped in the more acidic mitochondrial intermembrane space. This hypothesis is currently under further investigation.

### A new class of mitochondrial inhibitors

The proposed (H)CQ activity is a consequence in part of its protonatable nitrogens, which suggests that structurally related molecules might work in a similar fashion. To test this hypothesis (H)CQ activity was first optimized based on previous results. H293 cells in DMEM media plus FBS were allowed to attach overnight, then switched to L-15 galactose media and treated with the indicated CQ or HCQ concentrations for 16hrs (Figure 4A and B). Under these conditions oxidative phosphorylation was inhibited at much lower drug concentrations (compare to Figure 2D), consistent with time dependent (H)CQ accumulation in mitochondria. The addition of glucose largely overcame inhibition except at the highest drug concentration, suggesting that given enough time glycolysis may eventually be compromised (i.e. as in Figure 1A).

**Figure 4:**
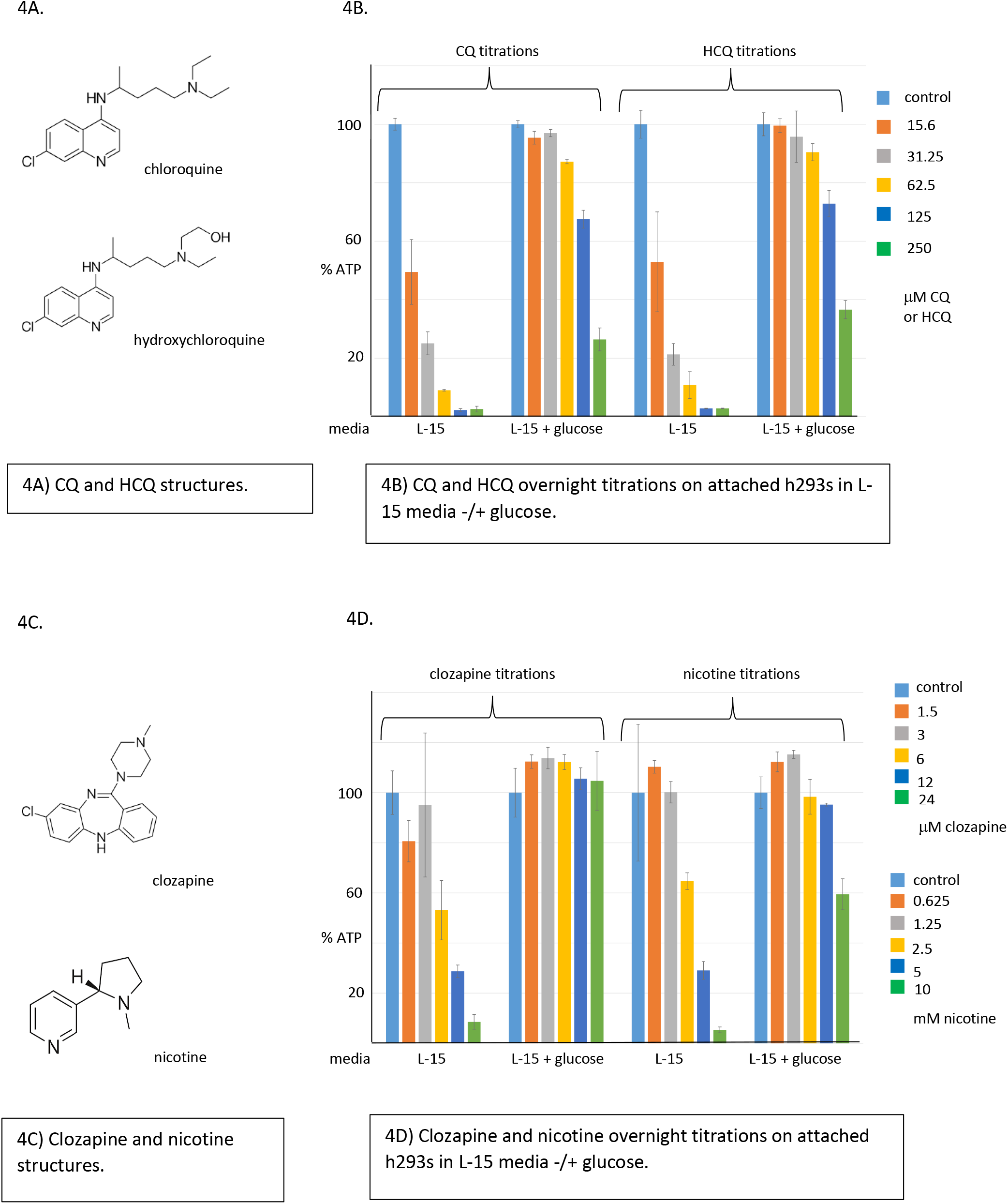
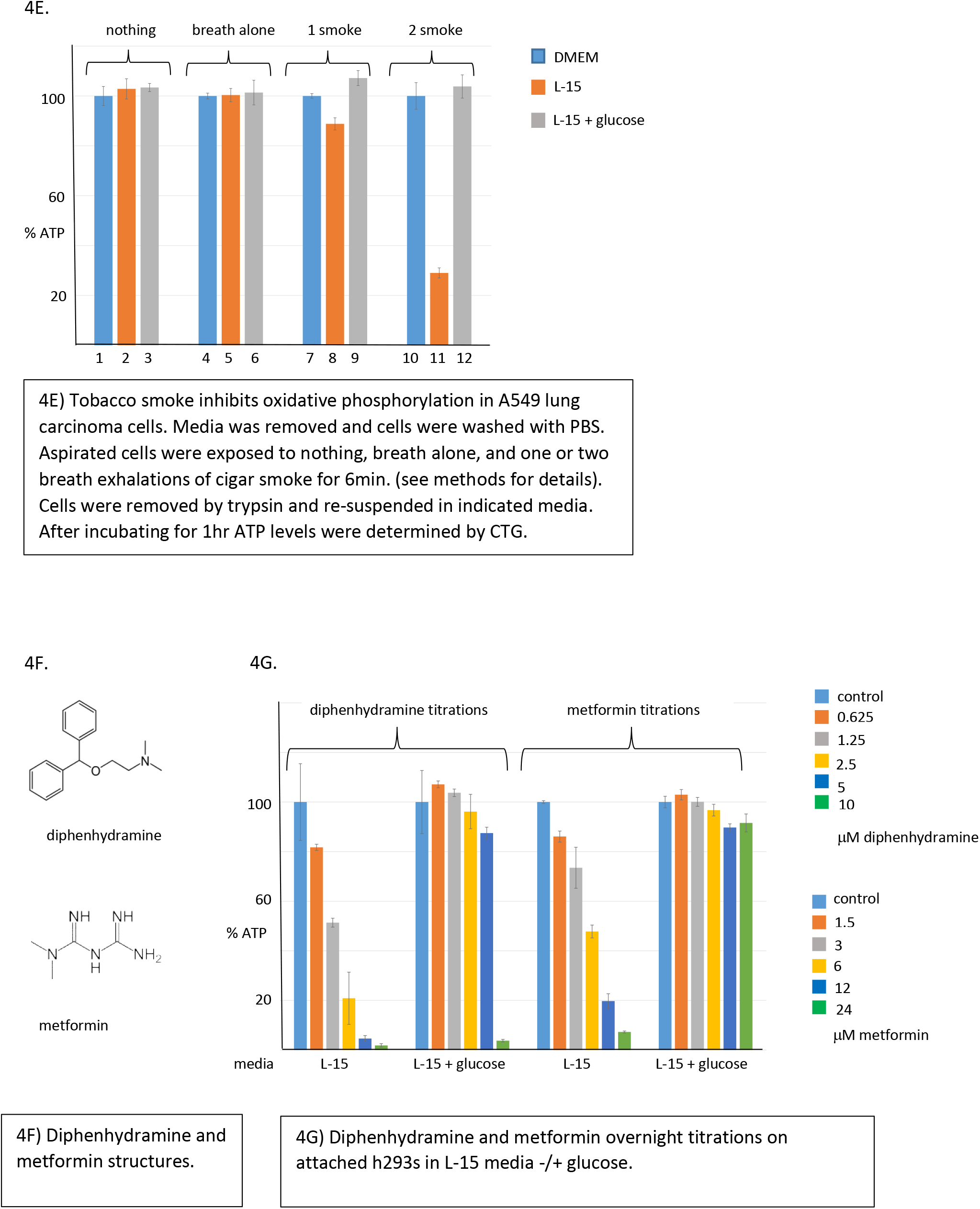
Molecules structurally similar to (H)CQ also target mitochondria.

Clozapine and nicotine were evaluated next (Figure 4C). Clozapine is an anti-psychotic medication that was identified as a potential COVID-19 therapeutic, and nicotine is of interest because epidemiological studies suggesting smokers may be protected from COVID-1.^18,19^ Both compounds preferentially decreased ATP levels when cells were forced to use oxidative phosphorylation, while glucose overcame this inhibition (Figure 4D). Note, however, that much higher nicotine concentrations were required. Attempts to enhance sensitivity by increasing the media pH (to favor generation of the free base) or by using different cells lines (e.g. A549 lung carcinoma cells) were unsuccessful (data not shown). Given that nicotine delivery to smokers is by inhalation, the effect of directly exposing cells to tobacco smoke was examined. ATP levels remained normal when smoke-treated cells were cultured in DMEM media that supports both glycolysis and oxidative phosphorylation. Remarkably, however, just 6min. smoke exposure dramatically decreased ATP levels when cells were forced to use the TCA/ETC pathway by culturing in L-15 galactose media (Figure 4E). Adding back glucose overcame inhibition by activating glycolysis, indicating that cell viability was not being affected. These results suggest that either nicotine vaporization is essential for efficient mitochondrial targeting, or some other component in tobacco smoke is responsible. These possibilities are currently under investigation., Based on the above results it was possible to identify additional compounds that would be expected to target mitochondria. Diphenhydramine, a common allergy medicine, was examined because it has two ring structures and contains a protonatable nitrogen attached to two methyl groups, similar to CQ (Figure 4F). Diphenhydramine had no effect when both metabolic pathways were available, but decreased ATP levels when cells were forced to use oxidative phosphorylation (Figure 4G). Conversely, the well-known diabetes drug metformin lacks ring constituents but does contain a nitrogen with two methyl groups (Figure 4F). It also specifically blocked ATP production by oxidative phosphorylation (Figure 4G), consistent with previous results suggesting it targets this pathway.^20^ In contrast, the potential COVID-19 therapeutic famotidine also has protonatable nitrogens, but completely failed to inhibit ATP synthesis (data not shown).^21^ Thus, entering cells and becoming protonated in the mitochondria appears to require additional structural elements that need to be identified.

## Discussion

Data suggest (H)CQ and related compounds are protonated and accumulate in mitochondria, which can compromise the electrochemical proton gradient needed to drive ATP synthesis. Given that in some cases (H)CQ appear to act prophylactically/therapeutically against COVID-19, their ability to sequester these protons raises the interesting possibility mitochondria are the initial target of SARS-Co-V2 infection. In this model (Figure 5) virus enters the cell via well-known mechanisms and forms an endosome (step 1).^5^ The endosome then fuses with a mitochondrion (step 2, which would provide protection from host defenses and a ready source of protons to release the viral genome (steps 3-5). This genome could exit and initiate virus production in the cytoplasm, or remain in the safe confines of the mitochondrion and utilize its ribosomes to produce more viral particles (step 6).^22^ The latter situation would eventually deplete protons and inhibit ATP synthesis, at which point virus could infect other mitochondria and repeat the process (steps 3-6). Once sufficient mitochondria are compromised and cellular ATP levels start to fall, glycolysis would be upregulated to compensate (step 7). The resulting lactic acid production (step 8) could lower cytosolic pH and create a more hospitable environment for extensive virus production in the cytoplasm (steps 9-13). In this model the initial mitochondrial infections are envisioned as a latent stage (relatively asymptomatic), and that of the cytoplasm as acute (extensive viral proliferation, viral particle release, and host cell damage).

**Figure 5:**
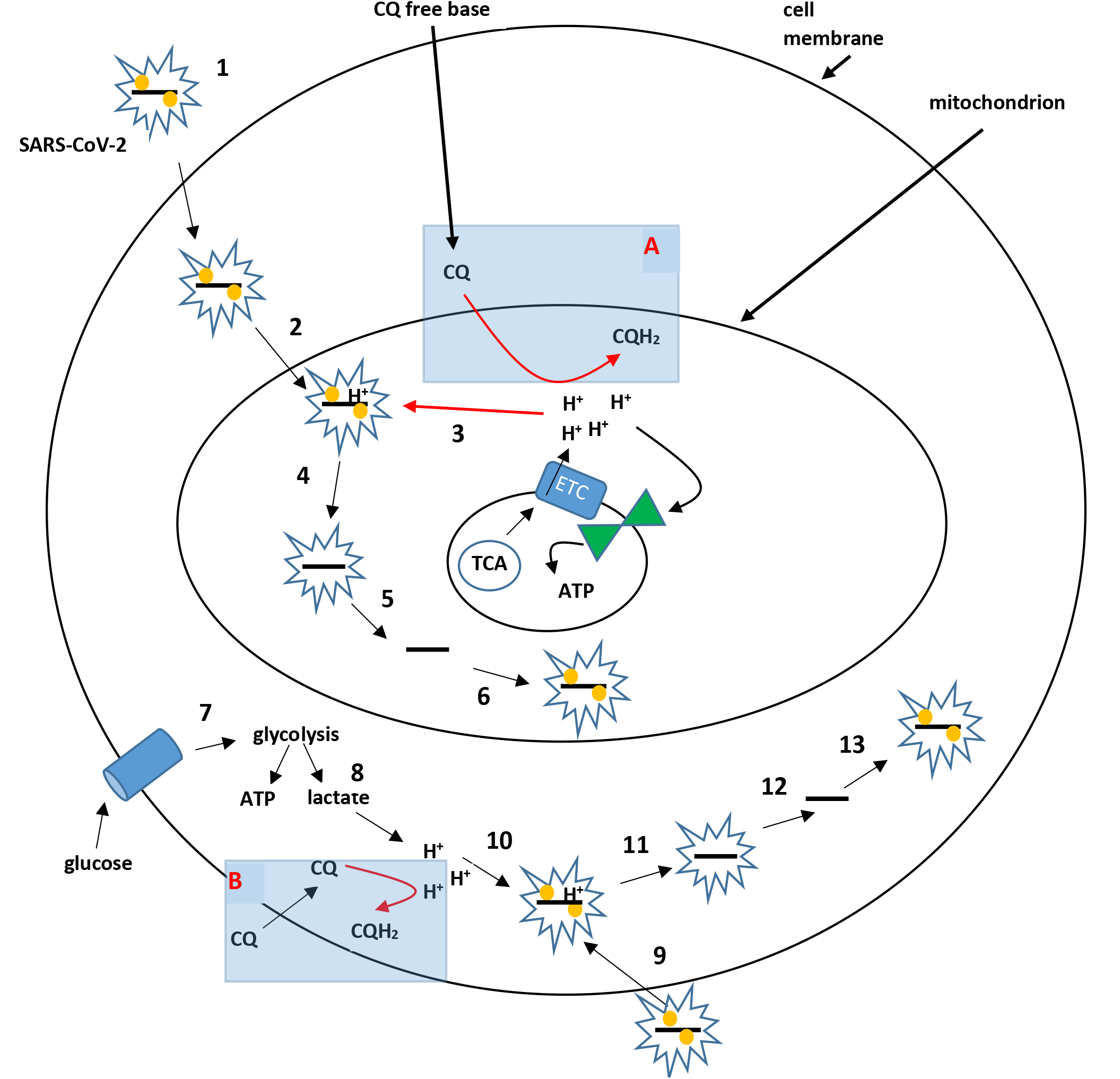
Model of SARS-CoV-2 infection and possible CQ/HCQ anti-viral activities.

Plasma membrane and mitochondrion of a host cell are depicted. Steps 1-5 propose the initial SARS-CoV-2 infection targets the mitochondrial intermembrane space, which hides the virus from host defenses and provides protons needed for genome release. Additional virus replication in the mitochondrion further depletes protons, which eventually inhibits ATP synthase activity (red hourglass) and forces the cell to upregulate glycolysis to produce ATP (steps 7 and 8). The glycolytic by-product lactic acid decreases cytoplasmic pH, which creates a cytosolic environment for increased viral production (steps 9-12). CQ and related molecules could act prophylactically and reduce the probability of initial infection by sequestering protons in the mitochondrial intermembrane space, thereby making this environment less hospitable for the virus (box A). If initial infection occurs, viral replication and expansion through the mitochondrial population could still be inhibited as depicted in box A. After oxidative phosphorylation is compromised and glycolysis upregulated, cytoplasmic acidification could lead to CQ protonation and inhibition of viral replication in this compartment (box B).

Based on the above model CQ and related compounds act prophylactically by decreasing acidity of the intermembrane space (box A) so that it is less conducive to viral infection (steps 1-3). This view is consistent with clinical preventative regimes employing long term low dosing of (H)CQ, which in conjunction with its long half-life allow slow mitochondrial accumulation at levels that do not severely compromise ATP production.^9^ Preliminary results indicate a component of tobacco smoke might function in this manner, which could explain lower infection rates for smokers.^19^ Even after initial infection (H)CQ could still act beneficially by slowing viral propagation through the mitochondria population. Proper dosing would be essential, however, because too much could actually inhibit oxidative phosphorylation and hasten the transition to cytoplasmic viral production. The well-known cardiotoxicity of CQ may be due to just such a phenomenon, as heart muscle is highly dependent on beta oxidation of fatty acids in mitochondria to produce ATP.^23^ Measuring blood lactate levels might be informative in this regard as a way to more precisely determine the CQ dose achieving maximal mitochondrial protection without upregulating glycolysis. Alternatively, a structurally related molecule (e.g. diphenhydramine or metformin) could show greater efficacy and reduced side effects compared to (H)CQ., As the disease progresses and glycolysis is upregulated, CQ could conceivably still act therapeutically by sequestering cytoplasmic protons and slowing viral replication (box B). However, its efficacy probably decreases dramatically at later stages of COVID-19 because: 1) Mitochondrial targets are already compromised, and 2) Tissue acidification from lactic acid excretion may protonate CQ and prevent cell entry. These ideas are also consistent with data from clinical studies indicating early CQ treatment yields better outcomes.^9^ A different approach at this stage, based upon the idea that infected cells are more dependent on glycolysis, might be therapeutic use of glycolytic inhibitors like 2-DG.^24^ They could perhaps be used alone or in combination with an (H)CQ-like molecule to cause apoptosis of infected cells by specifically decreasing their ATP levels. Uninfected cells should be resistant to such a treatment because the oxidative phosphorylation pathway is still functional and can metabolize amino acids., The proposed mitochondrial involvement in COVID-19 helps explain some curious features of the disease, such as its relatively asymptomatic effects on adolescents.^25^ The mitochondrial pool of young people adapt to increased energy requirements or partial inhibition by generating more mitochondria, which could extend the latent period of infection, perhaps indefinitely.^26^ Unfortunately, as this protective response declines over time the probability of disease progression to the acute phase increases. Dysregulation of metabolism might also help explain why obesity and diabetes correlate with more severe disease.^25,27^ Both these states exhibit higher blood glucose levels, which would be advantageous for infected cells dependent on glycolysis since it would help them maintain adequate ATP levels and re-direct carbon into viral biomass production. An additional adverse consequence of upregulating glycolysis is that the different metabolic byproducts (e.g. lactate) likely enter blood vessels, where they could disrupt the complex regulatory events controlling oxygen binding to hemoglobin.^28^, The SARS-CoV-2 virus causing COVID-19 has emerged as a serious and ongoing global threat. A biochemical analysis of (H)CQ mechanism of action in tissue culture cells suggests these and related drugs work in part by becoming protonated in the mitochondrial intermembrane space. This mechanism in turn raises the intriguing possibility that SARS-CoV-2 first infects and eventually compromises mitochondria, which preps the cytoplasm for additional virus replication and release by upregulating glycolysis and reducing cytosolic pH. This model, while speculative, is consistent with much of the biochemical and physiological data, provides ample opportunities for testing, and identifies novel approaches for therapeutic prevention and treatment of COVID-19.

## Materials and Methods

### Drugs

All drugs were obtained from Sigma except diphenhydramine (purchased commercially as 25mg allergy tablets) and famotidine (purchased commercially as 10mg acid controller tablets). Most compounds were dissolved in ultrapure water except quinine, which was re-suspended in 100% ethanol. Rotenone was prepared as a concentrated solution in 100% DMSO, then diluted to a working stock (125μM) in water (final concentration in reactions was 2.5μM). 2DG was used at a final concentration of 20mM. Diphenhydramine and famotidine were crushed by mortar and pestle, then re-suspended in H_2_0 and rotated at room temperature for 1hr. Insoluble material (e.g. cellulose) was removed by centrifugation. Drug presence was confirmed by UV/Vis analysis. Tobacco smoke was generated from commercially available Backwoods brand original cigars containing nicotine.

### Cell Culture

The following immortalized cell lines were obtained from the American Tissue Culture Collection: H293 kidney epithelial cells, HDFs (human diploid fibroblasts immortalized by telomerase expression), and U549 lung carcinoma cells. Cells were maintained in DMEM media (FisherScientific) supplemented with 10% fetal bovine serum (FBS; Atlas Biologicals, Fort Collins, CO), and penn/strep. It contained high glucose (25mM) and amino acids as carbon sources. Cultures were maintained in a 37°C water jacketed incubator with 5% CO_2_. When needed cells were removed from the plate using trypsin. It is important to note that FBS and many commercial preparations of trypsin contain glucose. After removal from the plate cells were gently pelleted by centrifugation (2K/4min. at room temperature), aspirated, and re-suspended in the desired media. Unless noted otherwise FBS was omitted because experiments showed it was not required for these short term metabolic assays (data not shown). Other medias used were DMEM no glucose and L-15, which contains amino acids but replaces glucose with 5mM galactose (both were from FisherScientific). L-15 also lacks sodium bicarbonate (NaHCO_3_), which was added to the same concentration as in DMEM (44mM) in all experiments except Supl. Fig. 3. Its presence is essential to buffer the media and prevent acidification in the CO_2_ incubator. A DMEM type media lacking sodium bicarbonate was also generated using individual components (FisherScientific and Sigma) at the concentrations listed in commercial preparations. This media was used for Figure 3C, where its pH was varied by the addition of 500mM Hepes pH 7-10 (25mM final concentration).

### Viability and ATP Assays

The CellTiterBlue (CTB) assay was obtained from Promega. It measures cell viability based on the ability of living cells to convert resazurin into the fluorescent resorufin. CellTiterGlo (CTG) was also from Promega. It lyses cells to release their ATP, which is used by a luciferase enzyme to produce light that is proportional to the ATP levels.

### Inhibition assays

In general, short term immediate effects of the indicated drugs on ATP levels were analyzed by removing growing cells from a p100mm plate using trypsin, washing with PBS, and re-suspending in the indicated media. After counting using a hemocytometer, 25-50k cells in 100μL media were distributed in a 96 well white flat bottom, non-treated polystyrene assay plate. In some cases media was supplemented by addition of glucose (20mM final). The plate was typically pre-incubated for 30min at 37°C in a CO_2_ incubator to adapt to the media. Drugs where then added in the order and at the concentration indicated for each figure, shaken (700RPM/10sec), and returned to the incubator., Incubation times were generally between 1 and 2hrs so that direct and immediate effects on ATP production could be evaluated. For the time course in Fig. 3B, cells were re-suspended in L-15 galactose media, treated with 250μM CQ and then incubated at 37°C 5% CO_2_ for the indicated times, followed by measurement of ATP levels using CTG. Untreated cells served as a control for each time point. ATP levels were determined by adding 10μL CTG reagent (Promega) directly to the wells followed by 5min. shaking at 700RPM. Light emitted by the ATP dependent luciferase was quantitated using a photoluminometer (BioTek Cytation 5). In most cases samples were run in duplicate or triplicate and the standard deviation calculated. Longer term effects on viability and ATP levels were analyzed by seeding cells in a tissue culture treated 96 well plate, allowing to attach overnight, then treating with indicated drug concentrations of drug for an additional 18hrs. To determine viability, 10μL CellTiterBlue resazurin reagent (Promega) was added and incubated for 2-4hr at 37°C in the CO_2_ incubator to allow conversion of resazurin to the fluorescent resorufin by viable cells. Fluorescence (ex^560^/em^590^) was measured on the BioTek Cytation 5. ATP levels were measure as described above using CTG. Because CellTiterBlue does not lyse cells, it was possible to add 10μL CellTiterGlo to the same wells, shake 5min., and determine ATP levels using CTG (Fig. 1). Luciferase activity alone (Supl. Fig. 1) was measured by adding 10μL CellTiterGlo reagent to 100μL DMEM media in the absence of cells. CQ (or drug of interest) was then added and the reaction initiated by the addition of 5μM ATP (final concentration). Luminescence was determined immediately and at later times as previously described. All drugs used in this study were tested and shown to not significantly inhibit the luciferase enzyme at the concentrations tested (data not shown).

### Fractionation

The published protocol of Clayton and Shadel was used with slight modifications.^29^ Briefly, 2.5×10^6^ h293 cells were re-suspended in 1ml L1-5 media without and with 20mM glucose. 1mM CQ in the same media but without cells served as a negative control, as did cells in the absence of CQ. Tubes were rotated 2hr. at room temperature to allow cellular entry of CQ. Cells were then pelleted by centrifugation (2K/4min. at room temperature), washed using L-15 media, and pelleted again. The cell pellet was re-suspended in 1ml ice-cold RSB hypotonic buffer and transferred to a cold 3ml dounce homogenizer. After allowing cells to swell for 10min. on ice, the cold B pestle was used to break them open (10 strokes)., Homogenization was confirmed by phase contrast microscopy. 0.7ml MS homogenization buffer was then added and the solution centrifuged at 1300g for 5min. at 4°C to remove nuclei (nuclear fraction). The supernatant was then centrifuged at 10,000g for 15min. generate a cytoplasmic fraction (supernatant) and a mitochondrial fraction (pellet). The nuclear and mitochondrial pellets were re-suspended in 750μL H_2_O, sonicated, and pelleted to remove insoluble material. The resulting cellular fractionations were analyzed directly for the presence of CQ by UV/Vis. Spectra were compared to that of purified CQ., For the red blood cell analysis, ~ 50μl blood was obtained from a finger prick and immediately added to PBS with 5mM EDTA to prevent coagulation. Cells were then pelleted and re-suspended in DMEM containing 5mM glucose (physiological concentration). After 2hr. incubation with 1mM CQ at room temperature while rotating, cells were pelleted, washed in PBS, and lysed in 750μL H_2_O with vortexing. After centrifugation to remove insoluble material, the supernatant was analyzed by UV/Vis spectroscopy. CQ in the absence of cells served as negative control.

### Smoke exposure

Four p60mm plates of attached A549 lung carcinoma cells were aspirated, washed with PBS, and aspirated again. Control plates were untreated and exposed to breath exhalation in the absence of smoke. Cigar smoke was blown into two experimental plates followed by rapid lid closure to trap the smoke. After 3min. at room temperature the lid was removed from one plate and smoke allowed to dissipate (labeled 1 smoke in Figure 4G). The other plate was exposed to another aliquot of cigar smoke followed by rapid lid closure (labeled 2 smoke). After 6 min. total incubation time cells were removed with trypsin, pelleted, and re-suspended in 500μL of the indicated media. 100μl aliquots were distributed in a white 96 well assay plate and incubated at 37°C in the CO_2_ incubator for 1hr. ATP levels were then analyzed using CellTiterGlo as described previously.

## Abbreviations

CQ: chloroquine
HCQ: hydroxychloroquine
2DG: 2-deoxy glucose
CTB: CellTiterBlue
CTG: CellTiterGlo
TCA: TriCarboxylic Acid cycle
ETC: Electron Transport Chain

## Acknowledgements

I would like to thank Dr. Syed Hussaini and Ian Mitchell for helpful suggestions, and The University of Tulsa for financial support.

## Author contributions

R.J.S. performed experiments and wrote the manuscript.

---

**Supl. Fig. 1:**
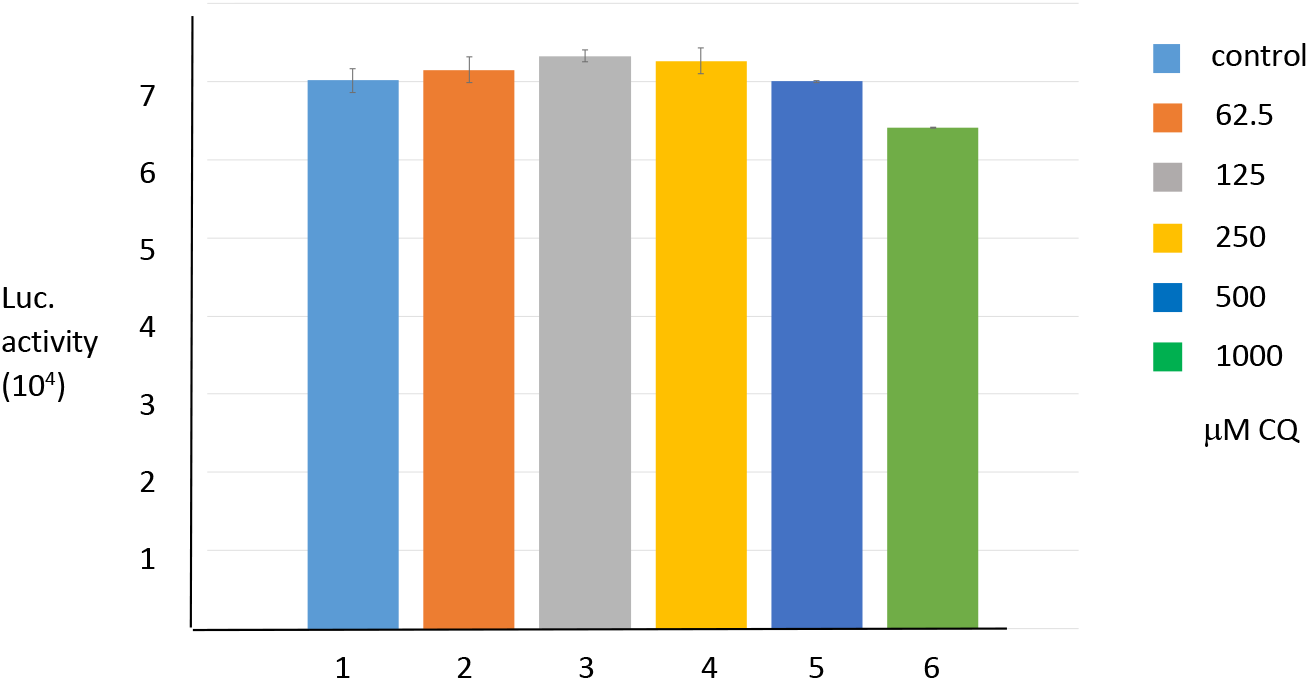
CQ does not inhibit the CTG luciferase reaction. CTG assay was performed in the absence of cells with exogenously added ATP and the indicated CQ concentrations.

**Supl. Fig. 2:**
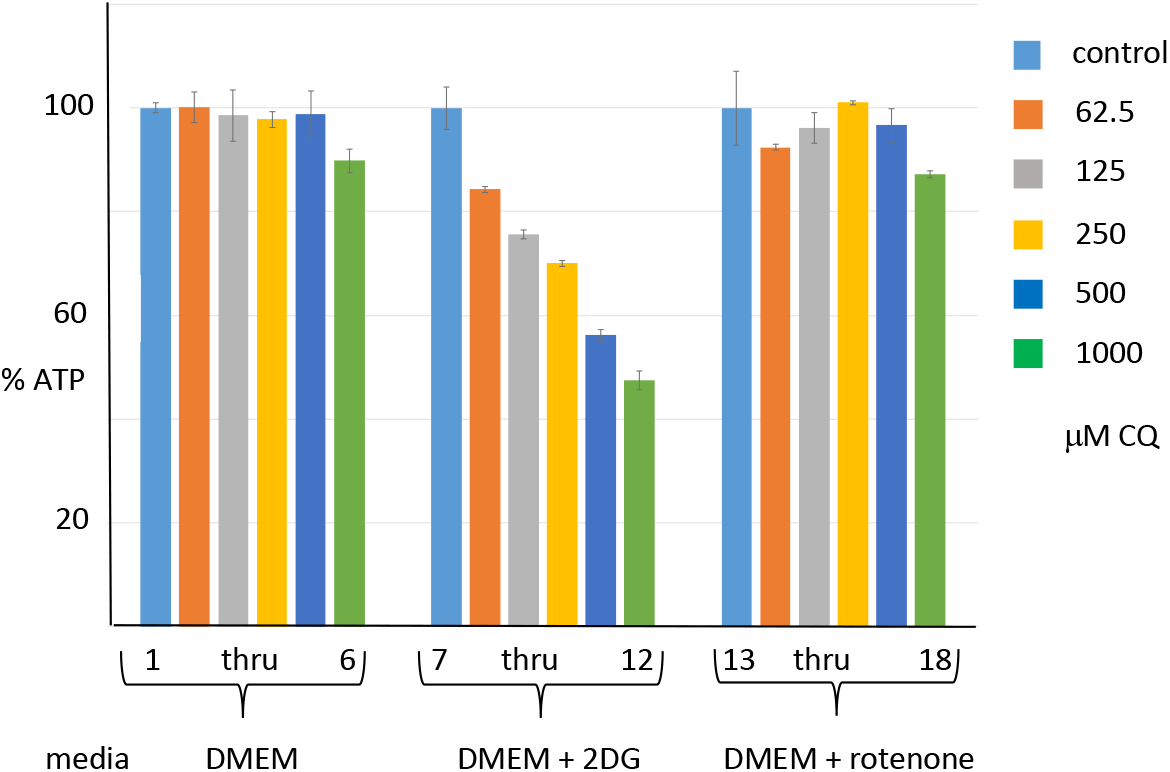
CQ preferentially decreases ATP levels in Human Diploid Fibroblasts preincubated with 2DG. HDFs suspended in DMEM were pretreated for 1hr with either 2DG (to force use of TCA/ETC) or rotenone (to force use of glycolysis). They were then exposed to the indicated concentrations of CQ for 2hr, followed by determination of ATP levels using CTG.

**Supl. Fig. 3:**
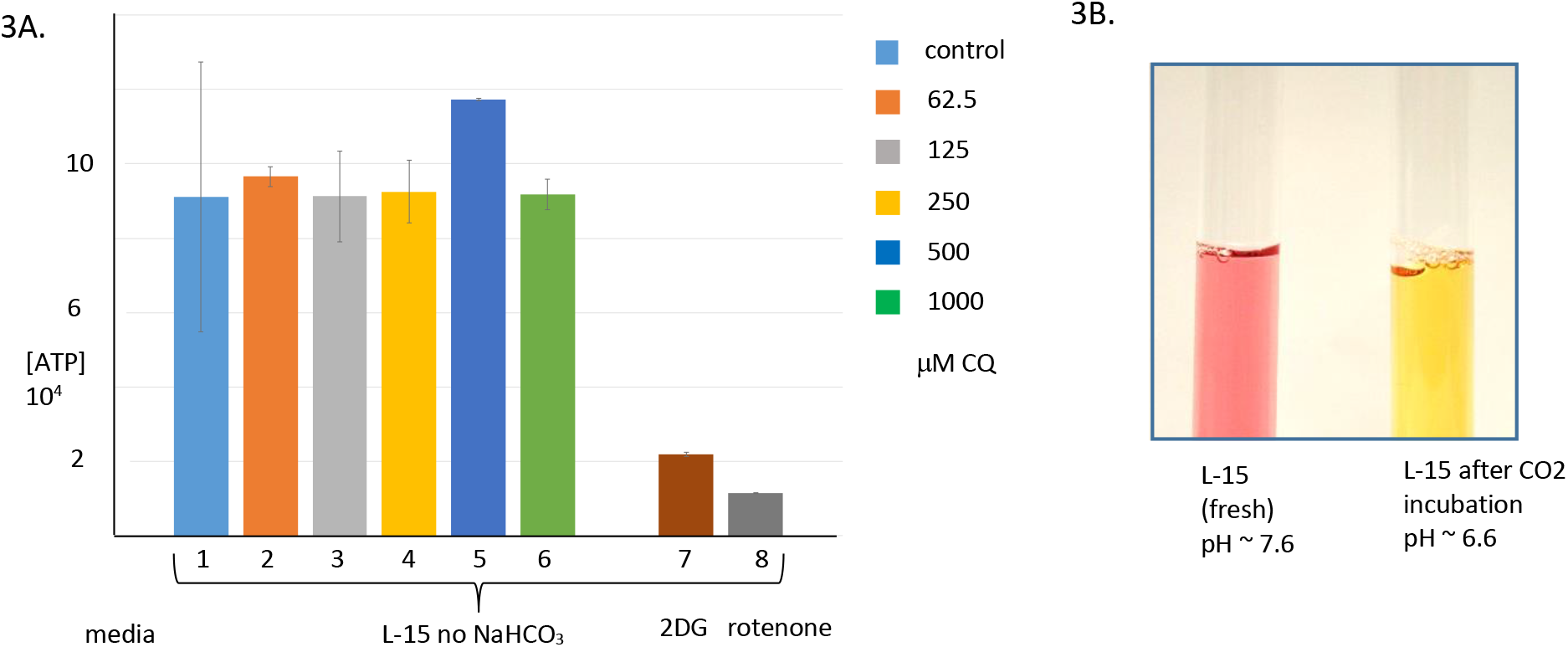
A) Media acidification blocks CQ inhibition. CQ titration (2hrs) on h293 cells cultured in L-15 media lacking NaHCO_3_ at 37°C in a CO_2_ incubator. 2DG (lane 7) and rotenone (lane 8) still inhibit ATP, indicating galactose is being metabolized. B) L-15 media color and pH change after 1hr CO_2_ incubation in the absence of NaHCO_3_.

**Supl. Fig. 4:**
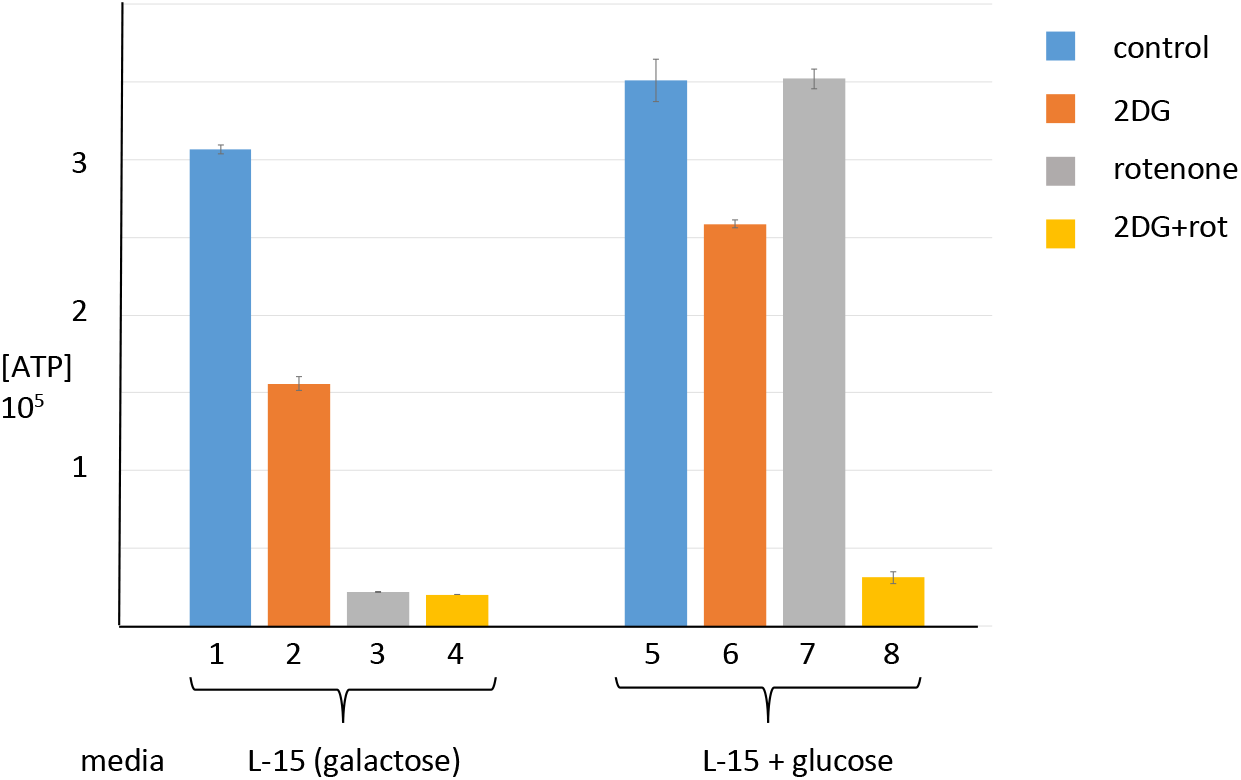
H293 cells in L-15 media containing galactose rather than glucose utilize oxidative phosphorylation. H293 cells refed with L-15 media were treated with 2DG, rotenone, or both inhibitors for 1hr to identify active metabolic pathways by measuring ATP levels using CTG (lanes 1-4). 20mM Glucose was added to L-15 media in lanes 5-8, which allowed glycolysis and reestablished the metabolic switch.

**Supl. Fig. 5:**
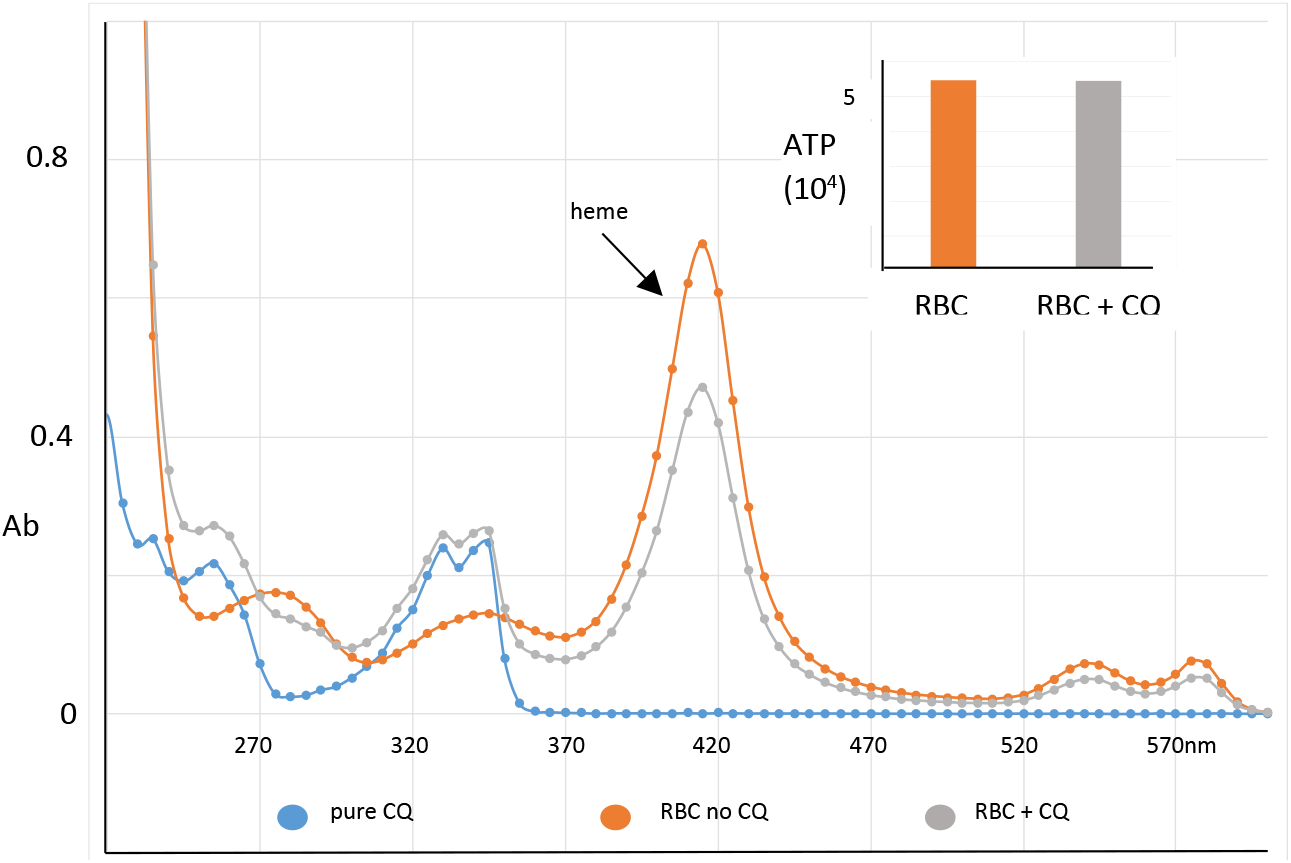
Red blood cells were pre-incubated with 1mM CQ for 1hr (orange line). After washing an aliquot was withdrawn to determine ATP levels using CTG (inset graph). Remaining cells were lysed by re-suspending in H2O and vortexing, followed by UV/Vis analysis of the supernatant. Pure CQ was analyzed to show its absorption peaks (blue line), while cells alone served as the negative control (grey line). The prominent heme peak is indicated by the arrow.

## References

1. Johns Hopkins University & Medicine. COVID-19 Dashboard by the Center for Systems Science and Engineering (CSSE) at Johns Hopkins University (JHU). https://coronavirus.jhu.edu/map.html.

2. Congressional Research Service. Global Economic Effects of COVID-19, https://fas.org/sgp/crs/row/R46270.pdf, 2020.

3. Walls AC, Park YJ, Tortorici MA, Wall A, McGuire AT, Veesler D. Structure, Function, and Antigenicity of the SARS-CoV-2 Spike Glycoprotein. Cell 181(2):281–292.e6. doi: 10.1016/j.cell.2020.02.058. PMID: 32155444, Epub 2020.

4. Alifano M, Alifano P, Forgez P, Iannelli A. Renin-angiotensin system at the heart of COVID-19 pandemic. Biochimie 174: 30–33, https://doi.org/10.1016/j.biochi.2020.04.008, Epub 2020.

5. Park SE. Epidemiology, Virology, and Clinical Features of Severe Acute Respiratory Syndrome - coronavirus-2 (SARS-CoV-2; Coronavirus Disease-19). Clin Exp Pediatr. 63(4):119–124. doi: 10.3345/cep.2020.00493, Epub 2020.

6. Malik YA. Properties of Coronavirus and SARS-CoV-2. Malays J Pathol. 42(1):3–11. PMID: 32342926, 2020.

7. Weiss SR, Leibowitz JL. Coronavirus pathogenesis. Adv Virus Res. 81: 85–164. doi: 10.1016/B978-0-12-385885-6.00009-2. PMID: 22094080, 2011.

8. Slater AF. Chloroquine: mechanism of drug action and resistance in Plasmodium falciparum. Pharmacol Ther. 57(2-3):203–35. doi: 10.1016/0163-7258(93)90056-j. PMID: 8361993, 1993.

9. Moore N. Chloroquine for COVID-19 Infection. Drug Saf. 1–2. doi: 10.1007/s40264-020-00933-4 PMCID: PMC7138193, Epub 2020.

10. Magagnoli J, Narendran S, Pereira F, Cummings T, Hardin JW, Sutton SS, Ambati J. Outcomes of hydroxychloroquine usage in United States veterans hospitalized with Covid-19. MedRxiv, 2020.04.16.20065920, DOI. 10.1101/2020.04.16.20065920, 2020.

11. Ferner RE, Aronson JK. Chloroquine and hydroxychloroquine in covid-19. BMJ. 369:m1432. doi: 10.1136/bmj.m1432. PMID: 32269046, 2020.

12. Fitch CD. Involvement of heme in the antimalarial action of chloroquine. Trans Am Clin Climatol Assoc. 109:97–105. PMID: 9601131, 1998.

13. Xue J, Moyer A, Peng B, Wu J, Hannafon BN, Ding WQ. Chloroquine is a zinc ionophore. PLoS One: 9(10):e109180. doi: 10.1371/journal.pone.0109180. eCollection. PMID: 25271834, 2014.

14. Schrezenmeier E, Dörner T. Mechanisms of action of hydroxychloroquine and chloroquine: implications for rheumatology. Nat Rev Rheumatol. (3):155–166. doi: 10.1038/s41584-020-0372-x. PMID: 32034323, Epub 2020.

15. Felsenstein S, Herbert JA, McNamara PS, Hedrich CM. COVID-19: Immunology and treatment options. Clin Immunol. 215:108448. doi: 10.1016/j.clim.2020.108448. PMID: 32353634, Epub 2020.

16. Guzik TJ, et.al. COVID-19 and the cardiovascular system: implications for risk assessment, diagnosis, and treatment options. Cardiovasc Res. cvaa106. doi: 10.1093/cvr/cvaa106. PMID: 32352535, Epub 2020.

17. Kase ET, Nikolić N, Bakke SS, Bogen KK, Aas V, G. Thoresen H, Rustan AC. Remodeling of Oxidative Energy Metabolism by Galactose Improves Glucose Handling and Metabolic Switching in Human Skeletal Muscle Cells. PLoS One 8(4): e59972. doi: 10.1371/journal.pone.0059972. PMCID: PMC3613401, 2013.

18. Remington G, Powell V. Clozapine and COVID-19. J Psychiatry Neurosci. 45(4):E1. doi:10.1503/jpn.2045301, 2020.

19. Vardavas CI, Nikitara K. COVID-19 and smoking: A systematic review of the evidence. Tob Induc Dis. 18: 20. doi: 10.18332/tid/119324. PMCID: PMC7083240, 2020.

20. Vial G, Detaille D, Guigas B. Role of Mitochondria in the Mechanism(s) of Action of Metformin. Front Endocrinol (Lausanne); 10:294. doi:10.3389/fendo.2019.00294, 2019.

21. Janowitz T, Gablenz E, Pattinson D, Wang TC, Conigliaro J, Tracey K, Tuveson D. Famotidine use and quantitative symptom tracking for COVID-19 in non-hospitalised patients: a case series. Gut. doi: 10.1136/gutjnl-2020-321852, Epub 2020.

22. Osellame LD, Blacker TS, Duchen MR. Cellular and molecular mechanisms of mitochondrial function. Best Pract Res Clin Endocrinol Metab. 26(6): 711–723. doi: 10.1016/j.beem.2012.05.003. PMCID:PMC3513836, 2012.

23. Kolwicz SC, Purohit S, Tian R. Cardiac Metabolism and its Interactions with Contraction, Growth, and Survival of Cardiomyocytes. Circulation Research:113 (5): 603–616. https://doi.org/10.1161/CIRCRESAHA.113.302095, 2013.

24. Pajak B, Siwiak E, Sołtyka M, Priebe A, Zieliński R, Fokt I, Ziemniak M, Jaśkiewicz A, Borowski R, Domoradzki T, Priebe W. 2-Deoxy-d-Glucose and Its Analogs: From Diagnostic to Therapeutic Agents. Int J Mol Sci. 21(1): 234. doi: 10.3390/ijms21010234 PMCID: PMC6982256 PMID: 31905745, 2020.

25. Dowd JB, Andriano L, Brazel DM, Rotondi V, Block P, Ding X, Liu Y, Mills MC. Demographic science aids in understanding the spread and fatality rates of COVID-19. Proceedings of the National Academy of Sciences 117 (18) 9696–9698; DOI: 10.1073/pnas.2004911117, 2020.

26. Sun N, Youle RJ, Finkel T. The Mitochondrial Basis of Aging. Mol Cell 61(5): 654–666. doi: 10.1016/j.molcel.2016.01.028, 2016.

27. Hussain A, Bhowmik B, Cristina do Vale Moreira N. COVID-19 and diabetes: Knowledge in progress. Diabetes Res Clin Pract. 162: 108142. doi: 10.1016/j.diabres.2020.108142. PMCID: PMC7144611, Epub 2020.

28. Kashani KB. Hypoxia in COVID-19: Sign of Severity or Cause for Poor Outcomes. Mayo Clin Proc. 95(6): 1094–1096. doi: 10.1016/j.mayocp.2020.04.021. PMCID: PMC7177114, Epub 2020.

29. Clayton DA, Shadel GS. Isolation of Mitochondria from Tissue Culture Cells. Cold Spring Harbor Protocols, doi:10.1101/pdb.prot080002, 2014.

